# Water-mediated interactions determine helix formation of peptides in open nanotubes

**DOI:** 10.1101/2020.11.11.378026

**Authors:** Dylan Suvlu, D. Thirumalai, Jayendran C. Rasaiah

## Abstract

Water-mediated interactions (WMIs) play diverse roles in molecular biology. They are particularly relevant in geometrically confined spaces such as the interior of the chaperonin, at the interface between ligands and their binding partners, and in the ribosome tunnel. Inspired in part by the geometry of the ribosome tunnel, we consider confinement effects on the stability of peptides. We describe results from replica exchange molecular dynamics simulations of a system containing a 23-alanine or 23-serine polypeptide confined to non-polar and polar nanotubes in the gas phase and when open to a water reservoir. We quantify the effect of water in determining the preferred conformational states of these polypeptides by calculating the difference in the solvation free energy for the helix and coil states in the open nanotube in the two phases. Our simulations reveal several possibilities. We find that nanoscopic confinement preferentially stabilizes the helical state of polypeptides with hydrophobic side chains, which is explained by the entropic stabilization mechanism proposed on the basis of polymer physics. Polypeptide chains with hydrophilic side chains can adopt helical structures within nanotubes, but helix formation is sensitive to the nature of the nanotube due to WMIs. We elaborate on the potential implications of our findings to the stability of peptides in the ribosome tunnel.

## Introduction

Understanding proteins in crowded and confining environments is important for knowledge of their function in the cell.^1–3^ For example, experiments suggest that the architecture of the ribosome promotes transient structure acquisition in newly synthesized proteins.^4? –12^ Theoretical studies predicted that the ribosome tunnel stabilizes the *α*-helix in proteins by destabilizing the coiled state through a reduction in entropy of the coiled state due to conformational restrictions from the confining walls.^13–15^ However, molecular dynamics simulations of a hydrophobic polypeptide confined to a periodically replicated carbon nanotube containing water showed that the coiled state is preferred in the nanotube,^16^ which appears to naively contradict the expectations based on polymer physics. ^13^ The cause of this surprising behavior was attributed to the presence of water which was confined within the nanotube.^17^ Removal of periodic replication of the nanotube and exposure to a water reservoir completely changes the picture, and the conditions for helix formation now occurs within a range of hydrophobicity and tube diameters similar to the diameters of the ribosome tunnel.^18^ Additional studies indicate a central role for water in contributing to conformational preferences of biomolecules in crowded and confining environments,^19–23^ and water has been receiving increasing attention for its role in biology. For example, the hydrophobic effect, studied in a series of pioneering papers initiated by Pratt,^24,25^ is thought to play a major role in protein folding.^26–29^ Further, water-mediated interactions (WMIs) have been demonstrated to play a role in protein aggregation and amyloid formation and amyloid polymorphism.^30,31^ It has even been speculated that water may be the universal solvent for life. ^32^ However, WMIs are not completely understood, and surprises^33,34^ and new perspectives continue to emerge^35^ largely because the variability in water-mediated interactions are dictated by the context. Therefore, WMIs continue to be an abiding topic of interest and characterizing their influence remains to be a fruitful enterprise.^36–39^

Conformational preferences of biomolecules in nanoscopic confinement are difficult to predict due to the presence of many competing interactions. For example, one must consider not only intramolecular interactions within the protein, but also intermolecular interactions with the confining surface and how both of these interactions are mediated by water within and outside a nanotube open to a reservoir. The latter water-mediated component of these interactions is inherently a many-body effect, thus making it difficult to describe theoretically. Further, water can behave in surprising ways in confined spaces^40–43^ leading to additional complexity. Here, we employ a model system containing a homopolymer with either a hydrophobic or hydrophilic amino acid confined to polar or non-polar nanotubes in the gas phase and in the solution phase when the nanotubes are open to a water reservoir. At equilibrium, the chemical potential of water within and outside the open nanotube in solution is identical. The model benefits from being simple enough to extract meaningful trends, but complex enough to potentially be relevant to biological systems. Using molecular dynamics simulations, we demonstrate that water-mediated interactions (WMIs) contribute to conformational preferences of polypeptides in nanotubes. We quantify WMIs in this context by calculating the difference in the solvation thermodynamics between the helix and coil states while confined to non-polar and polar nanotubes in the gas and solution phases. Our simulation results suggest that hydrophobic sequences preferentially form *α*-helices inside open nanotubes. The results presented here should be of interest to researchers studying water-mediated interactions and confinement effects on biomolecules.

## Methods

Our liquid water simulations followed a similar procedure to our previous work. ^18^ We performed replica exchange molecular dynamics simulations^44^ using GROMACS 5.1.2^45^ with the CHARMM36 force field.^46,47^ The nanotubes were 100 Å in length with diameters 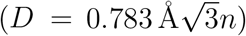 13.6, 14.9, 16.3, 17.6, and 18.9 Å. The 13.6 and 14.9 Å NT systems each contained 74 replicas, while the 16.3, 17.6, and 18.9 Å NT systems each contained 84 replicas. The replicas spanned the temperature range 280 K to 500 K with the temperature spacing between replicas determined by a procedure described elsewhere^48^ such that the exchange probability between replicas was at least 0.23 (±0.02). Exchange attempts between replicas were attempted every 1 ps. Errors were estimated with block averaging.^49^ The gas phase simulations followed a procedure used previously to study protein dynamics in the gas phase. ^50,51^ The number of replicas for each NT system in the gas phase was 30, spanning the temperature range 280 K to 600 K with temperature spacing such that the exchange probability between replicas was approximately 0.30 (±0.01). Additional details of the simulation methods can be found in the Supporting Information.

## Results and Discussion

To compute the difference in the solvation free energies between the helix (h) and coil (c) states, we constructed a thermodynamic cycle (Fig. 1). The solvation free energy of the helix is 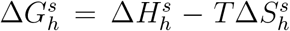 where 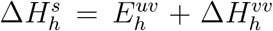 and 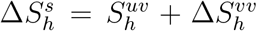.^52–55^ The solute (polypeptide + nanotube) is denoted by *u* and the solvent (water) is denoted by *v*. The quantity 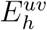 is the average interaction energy between the solute and solvent, and 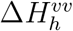 is the water reorganization energy upon solvation of the helical state.^52–55^ Further, the water reorganization energy is exactly compensated by the water reorganization entropy, 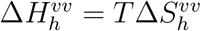.^52–55^ The quantity 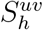 is the corresponding entropy associated with solute-solvent interaction energy fluctuations. ^52–55^ A similar expression holds for the coil. The difference in solvation free energies is,

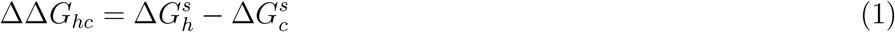

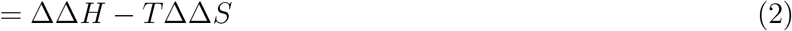

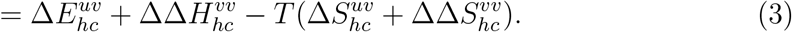

**Figure 1:**
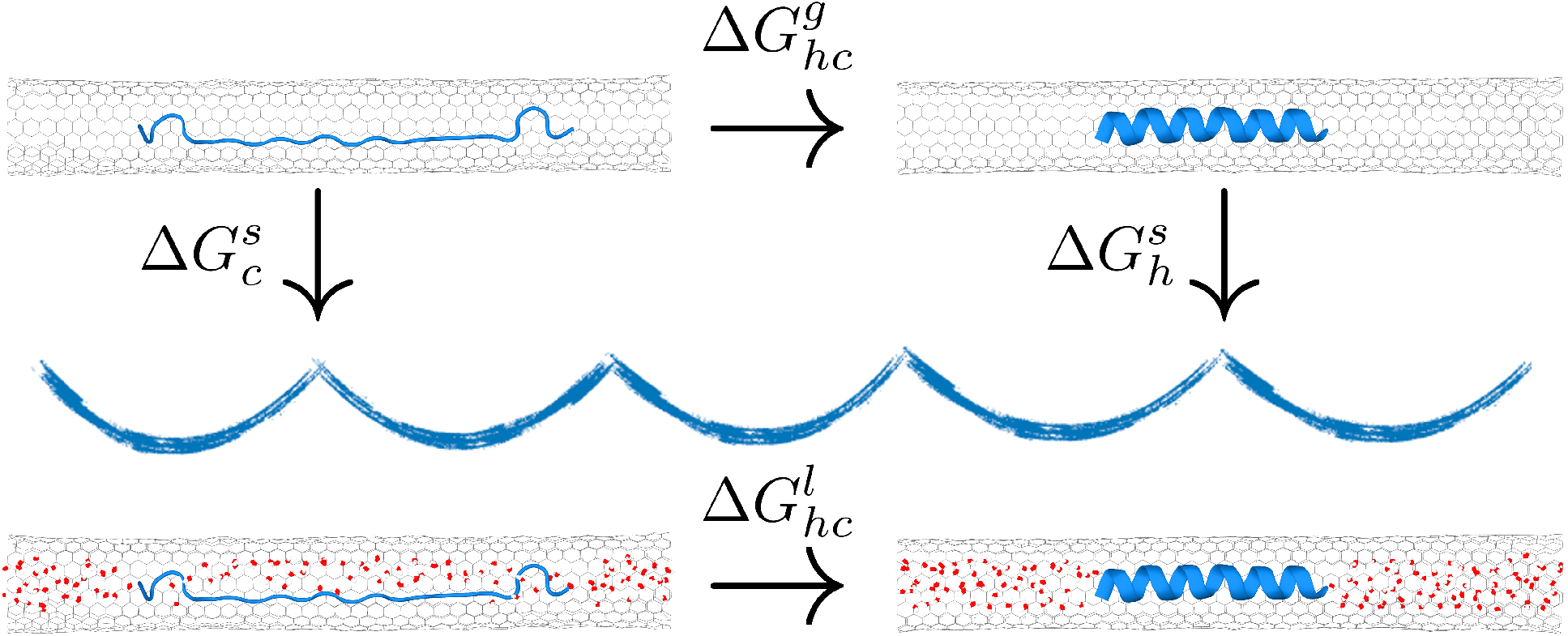
Thermodynamic cycle used in calculating ΔΔ*G_hc_*, the difference in solvation free energies between the helix and coil states in the gas and liquid phases.

Because the Gibbs free energy is a state function, the difference in solvation free energies can be computed through the difference in the free energies of the helix and coil states in the liquid 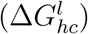 and gas phases 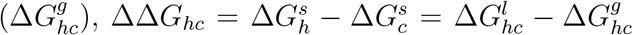. Both 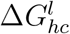 and 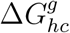 can be obtained from replica exchange molecular dynamics simulations between identical systems over a range of temperatures, from which we obtain an estimate of ΔΔ*G_hc_* and, therefore, insight into the water-mediated contributions to helix formation inside nanotubes open to a water reservoir.

We computed ΔΔ*G_hc_* for four systems. (1) A polypeptide (A_23_) with a non-polar side chain (–CH_3_) confined to a non-polar carbon nanotube (NPNT). (2) A polypeptide (S_23_) with a polar side chain (−CH_2_OH) confined to a non-polar carbon nanotube. (3) A_23_ confined to a polar boron nitride nanotube (PNT), and (4) S_23_ confined to a polar boron nitride nanotube.

The free energy of helix formation in the liquid and gas phases was computed as 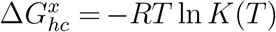 where the temperature dependent equilibrium ratio *K*(*T*) = *f_h_*(*T*)/(1 – *f_h_*(*T*)). The fraction of the polypeptide in the helix state *f_h_* is obtained through a Lifson-Roig formalism for helix-random coil transition of polypeptides as was done in our previous publication^18^

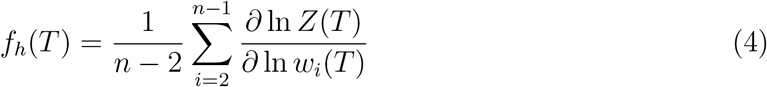

where *Z* is the partition function of the polypeptide, *n* is the number of amino acids, and *w_i_*(*T*) is a weight factor for each amino acid.^56^ Additional details about how the thermodynamics at 300 K were determined are provided in the Supporting Information.

Figure 2 displays the fraction helix of A_23_ and S_23_ confined to the NPNT and PNT with diameters 13.6 Å and 14.9 Å, in the liquid and gas phases as a function of temperature. Inside both the non-polar nanotube (NPNT) and polar nanotube (PNT), A_23_ has greater helix content in the presence of water compared to the gas phase. Conversely, at 300 K S_23_ forms a helix inside the NPNT in the gas phase but not in the presence of water. However, the helix content of S_23_ is reestablished in the PNT when in the presence of water. We elaborate on these observations by discussing the difference in free energies of helix and coil states in the gas and liquid phases ΔΔ*G_hc_* for the four systems.

**Figure 2:**
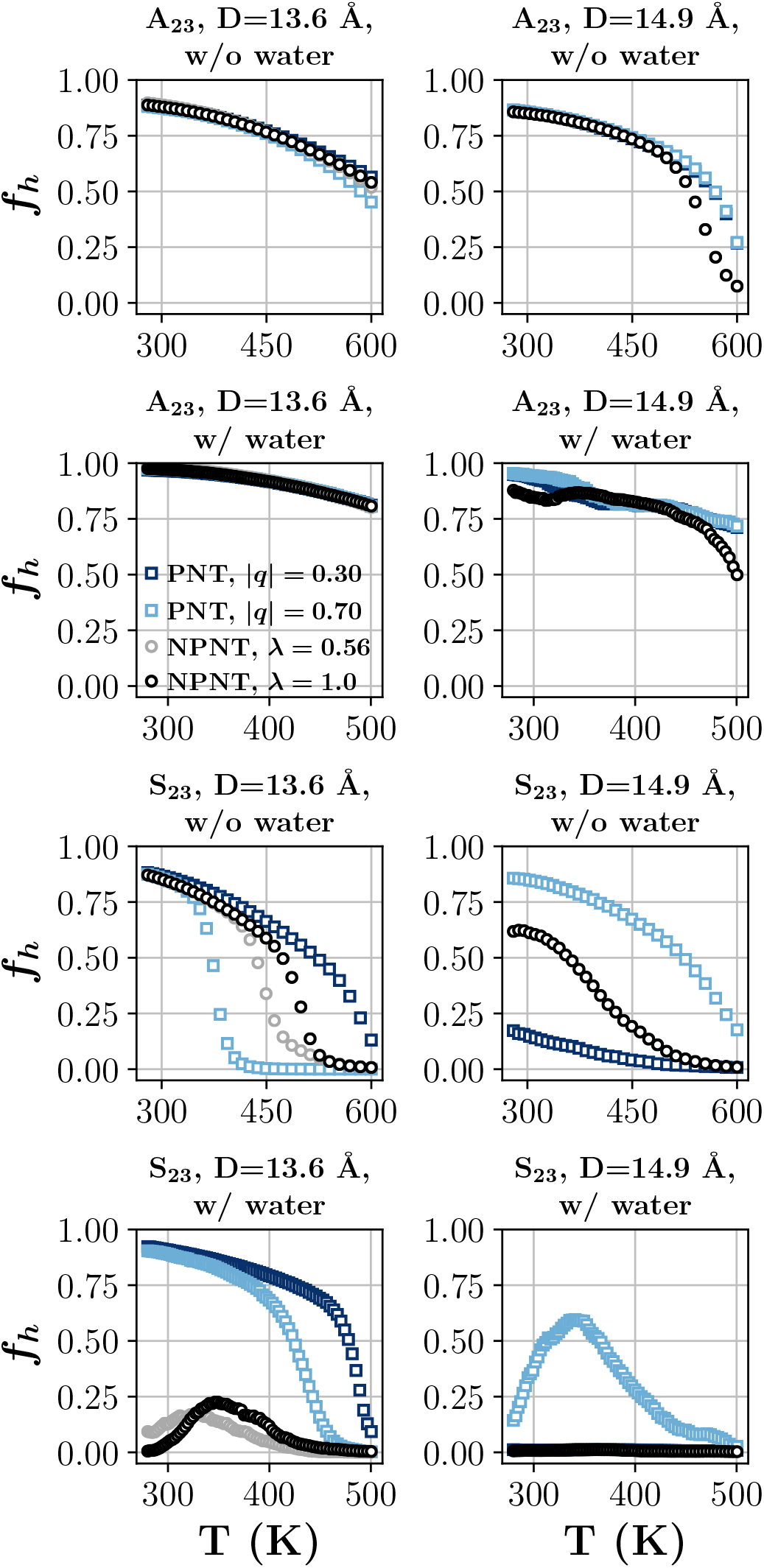
Fraction helix *f_h_* of polyalanine (A_23_) and polyserine (S_23_) as a function of temperature while confined inside the non-polar carbon nanotube (NPNT) and polar boron nitride nanotube (PNT) for diameters *D* = 13.6Å (left column) and *D* = 14.9 Å (right column) both with and without water.

### NPNT/A_23_

We calculated the free energy of helix formation in A_23_ inside a carbon nanotube in the gas phase and in liquid water (Fig. 3). Compared to the gas phase, the helix state of A_23_ in the presence of water has a lower free energy with ΔΔ*G_hc_* ≈ −4kJ/mol (Table 1). In other words, it is more favorable to solvate the A_23_ helix/nanotube system than the A_23_ coil/nanotube system. Table 1 suggests that this is an energetic rather than an entropic effect. Specifically, ΔΔ*H* < 0 while ΔΔ*S* < 0. However, upon closer inspection we will find that this must be driven by the entropy. In the coil state, A_23_ has many more hydrogen bonds with water than in the helix state. Therefore, the water-polypeptide interaction energy is expected to be more negative for the coil than the helix, 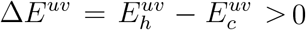. Consequently, the difference in water reorganization energy between the helix and the coil states must be negative, 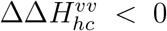. See the Supporting Information for further evidence of this. Accordingly, one might expect water reorganization energy to favor the helix state inside the nanotube, and provide greater helix stability for A_23_ in the liquid phase compared to the gas phase. However, the water reorganization energy and entropy exactly compensate, Δ*H^vv^* = *T*Δ*S^vv^*, as discussed above. Therefore, 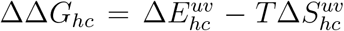. For ΔΔ*G_hc_* < 0, as is the case for A_23_ confined to the NPNT, the entropy associated with the fluctuations in the solute-solvent interaction energy must be positive, 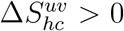. Thus, the greater helix stability conferred to the peptide in the liquid phase compared to the gas phase originates from the positive entropy change associated with the fluctuations in the water-solute interaction energy.

**Figure 3:**
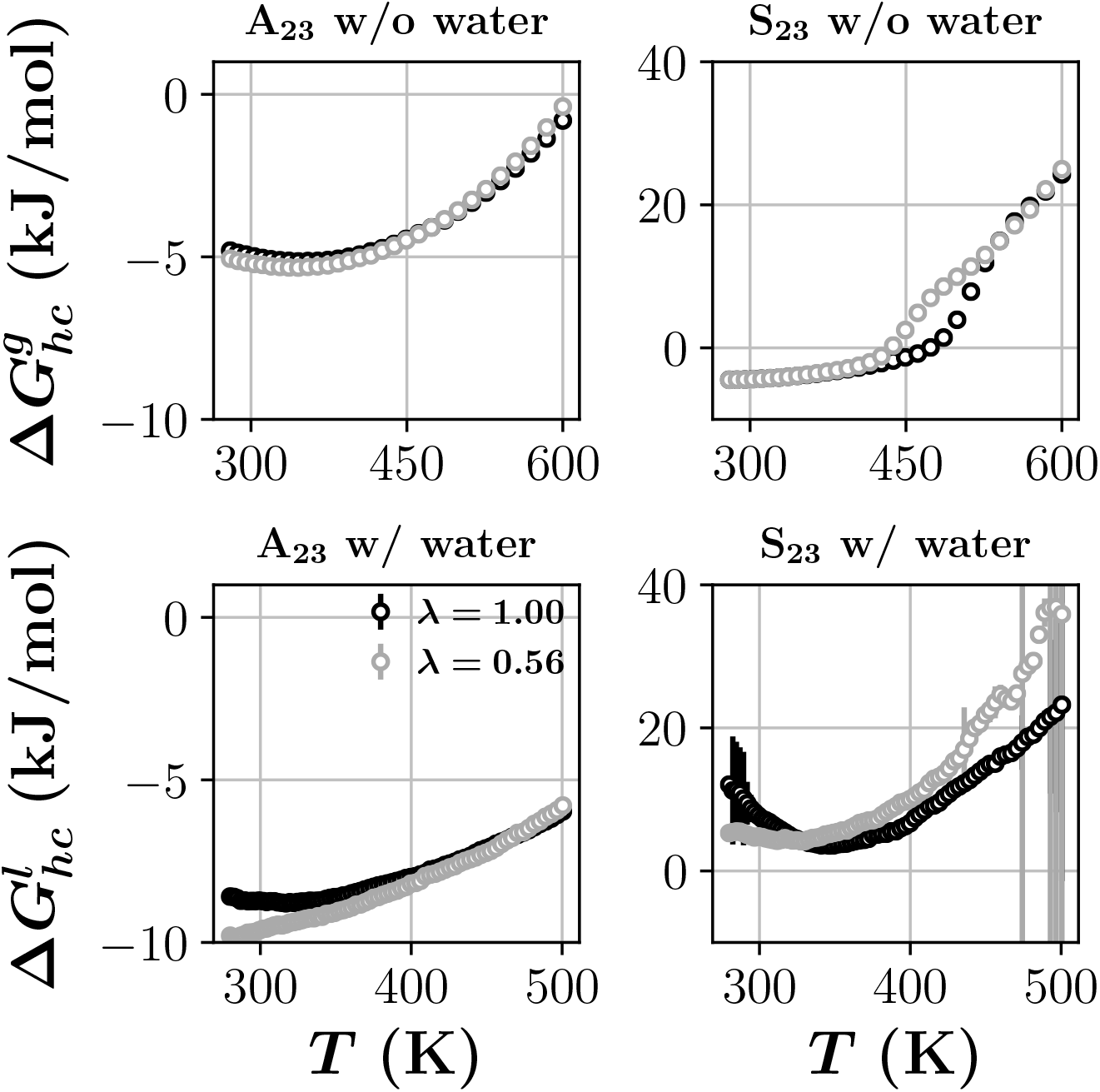
Free energy of helix formation in polyalanine (A_23_) and polyserine (S_23_) confined inside 13.6 Å non-polar carbon nanotubes both with and without water.The parameter λ scales the Lennard-Jones potential of the carbon nanotube.

**Table 1:** Differences in free energy, enthalpy, and entropy for the helix and coil states at 300 K for A_23_ and S_23_ confined to the non-polar carbon nanotube of diameter *D* = 13.6 Å.

| | $\Delta\Delta G_{hc}$ (kJ/mol) | $\Delta\Delta H$ (kJ/mol) | $\Delta\Delta S$ (J/mol) |
| --- | --- | --- | --- |
| A<sub>23</sub> |  |  |  |
| $\lambda = 1.00$ | $-3.79 \pm 0.25$ | $-4.42 \pm 0.18$ | $-2.1 \pm 0.6$ |
| $\lambda = 0.56$ | $-4.42 \pm 0.39$ | $-9.29 \pm 0.27$ | $-16.2 \pm 0.9$ |
| S<sub>23</sub> |  |  |  |
| $\lambda = 1.00$ | $11.1 \pm 2.3$ | $48.8 \pm 1.7$ | $125.6 \pm 5.2$ |
| $\lambda = 0.56$ | $9.2 \pm 1.5$ | $25.8 \pm 1.1$ | $55.3 \pm 3.4$ |

We also scaled the dispersion energy of the carbon nanotube with a parameter λ as in our previous study.^18^ The free energy of helix formation inside the NPNT in the gas phase is relatively insensitive to reductions in the dispersion energy. In the presence of water, however, the free energy of helix formation decreases by approximately 1 kJ/mol as λ decreases from 1.0 to 0.56. The increase in helix stability as the dispersion energy of the nanotube decreases suggests a dewetting phenomena contributes to helix formation in this case. The surface of the A_23_ helix is hydrophobic (Fig. 4) as is the inside surface of the NPNT. During helix formation, these two surfaces associate while simultaneously expelling water from the region between them into the reservoir. This dewetting phenomena becomes more likely as the dispersion energy of the nanotube decreases since there are fewer water molecules in the tube and relatively large water density fluctuations become more likely. ^27,57,58^

**Figure 4:**
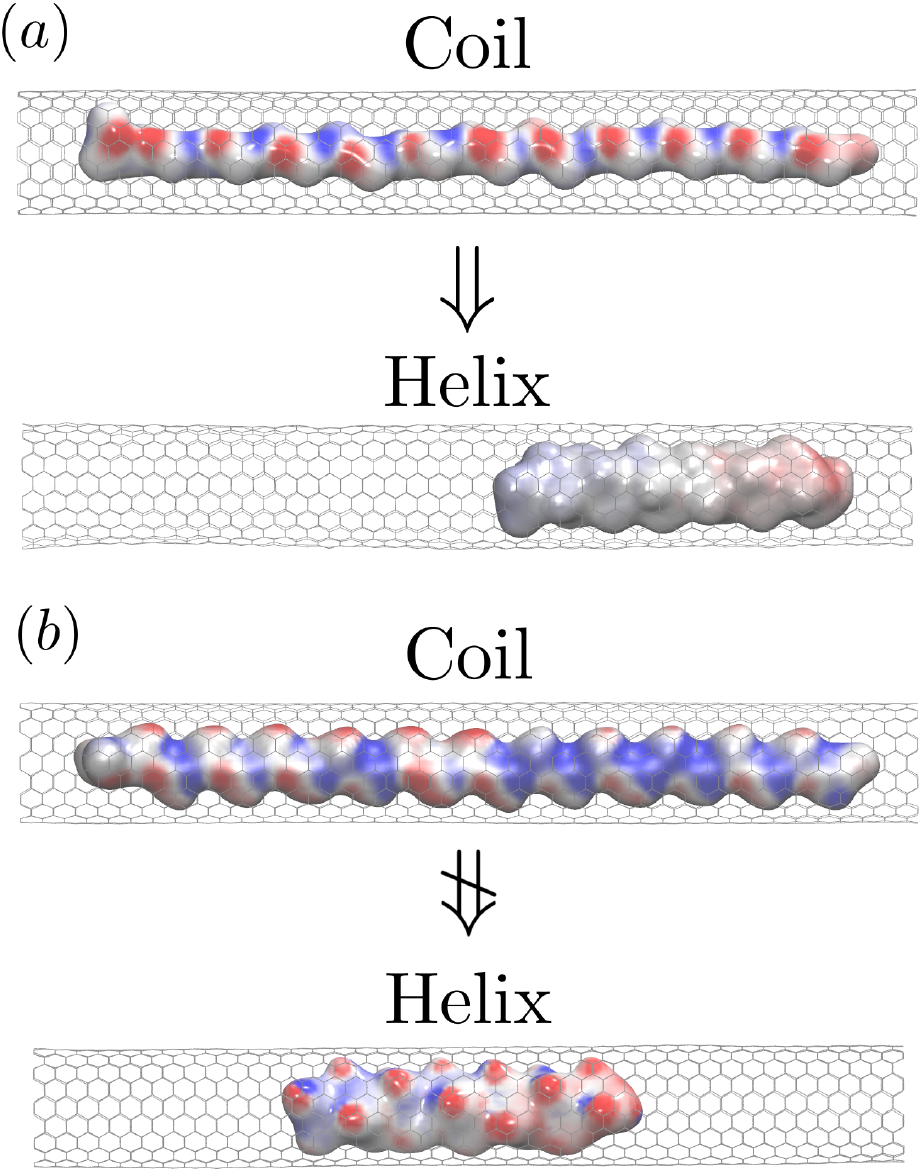
Electrostatic potential from the Poisson-Boltzmann equation for the coil and helix states for (a) A_23_ and (b) S_23_ inside the NPNT. Blue indicates positively polarized (-NH), red indicates negatively polarized (-CO and -OH), and gray indicates neutral or non-polar. Note the non-polar nature of the surface of the A_23_ helix, and the polar nature of the surface of the S_23_ helix.

### NPNT/S_23_

In contrast to A_23_, which has a hydrophobic side chain, S_23_ has a hydrophilic side chain (Fig. 4). Therefore, the water-mediated interactions between S_23_ and the nanotube are expected to be different from A_23_. Indeed, we observe helix formation in S_23_ in the NPNT in the gas phase at 300 K which is expected from polymer physics. However, we do not observe helix formation in S_23_ the presence of water inside the NPNT. This is further demonstrated by the negative change in free energy for helix formation at 300 K in the gas phase, but positive free energy change for helix formation in the presence of water (Fig. 3). The result is a positive difference in free energies between the helix and coil states for S_23_ (Table 1). In other words, solvation of the S_23_ coil/nanotube system is more favorable than the S_23_ helix/nanotube system when the nanotube has non-polar walls. The -OH functional group of S_23_ forms approximately 42 hydrogen bonds with water molecules inside the *D* = 13.6 Å nanotube (Fig. 5). To form a helix, S_23_ would have to break many of these hydrogen bonds because at *D* = 13.6 Å there is not enough room inside the nanotube to maintain a hydration layer around the helix (Fig. 4). Therefore, 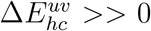 and ΔΔ*H* >> 0. Further evidence of this is displayed in the Supporting Information. Consequently, helix formation is not favored in S_23_ inside the NPNT. The release of the hydrogen bonded water from the serine hydroxyl group during helix formation would result in an increase in the entropy of the bound water due to the increase in translational freedom. This is reflected in the positive ΔΔ*S* in Table 1, but its magnitude is not large enough to compensate for the energy required to desolvate the peptide side chain. In this case, it could be expected that the helix content in S_23_ might increase with temperature as water’s tendency to form hydrogen bonds decreases with increasing temperature. This is observed in the temperature dependence of the helix content in S_23_ in the NPNT, with a maximum in *f_h_* observed at approximately 350 K before approaching zero for *T* > 350 K (Fig. 2).

**Figure 5:**
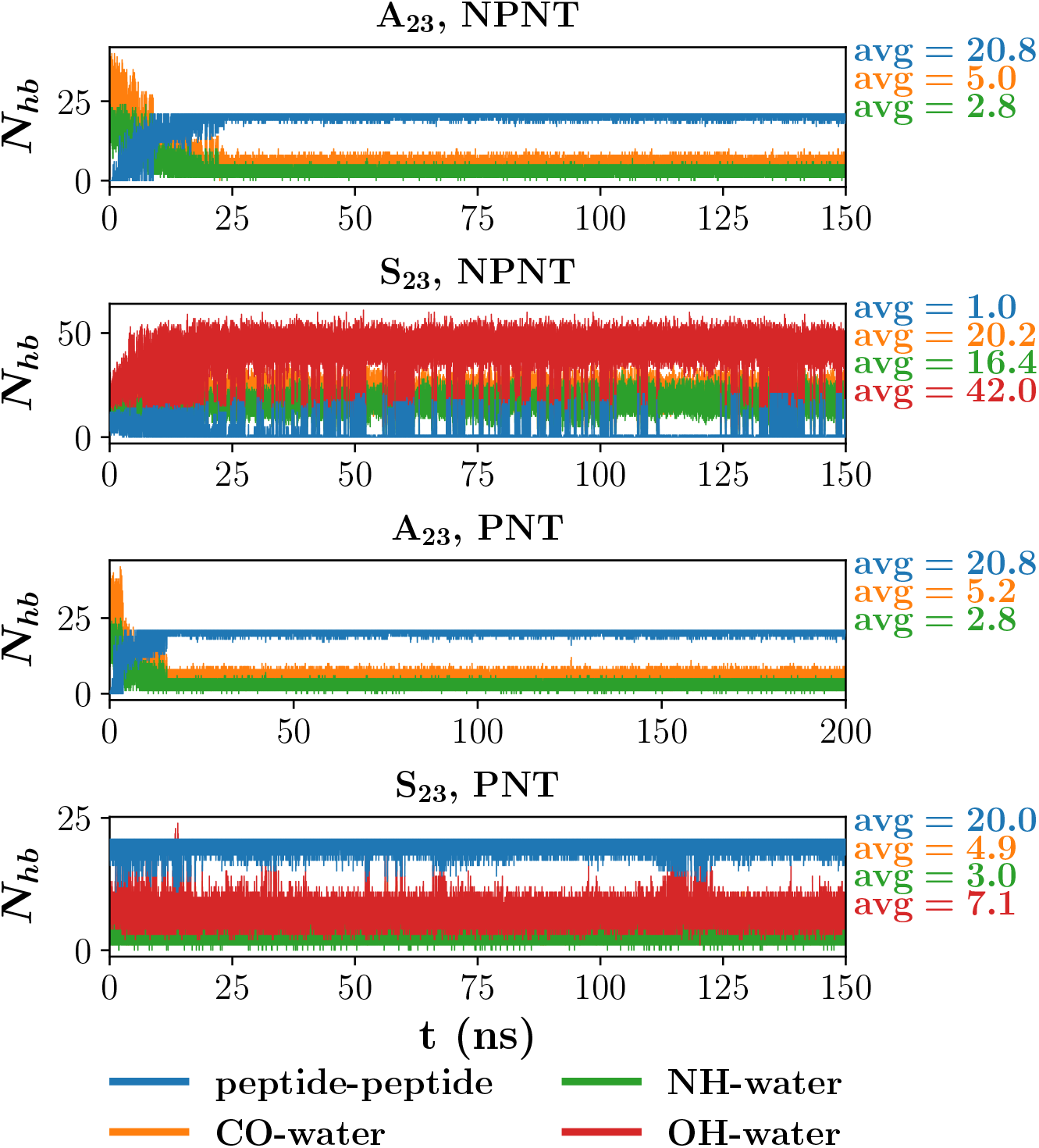
Number of peptide-peptide and peptide-water hydrogen bonds (*N_hb_*) as a function of time at 300K for A_23_ and S_23_ confined to the NPNT (λ = 1.00) and PNT (|*q*| = 0.30). The displayed averages are calculated from the last 50 ns of the simulations.

**Figure 6:**
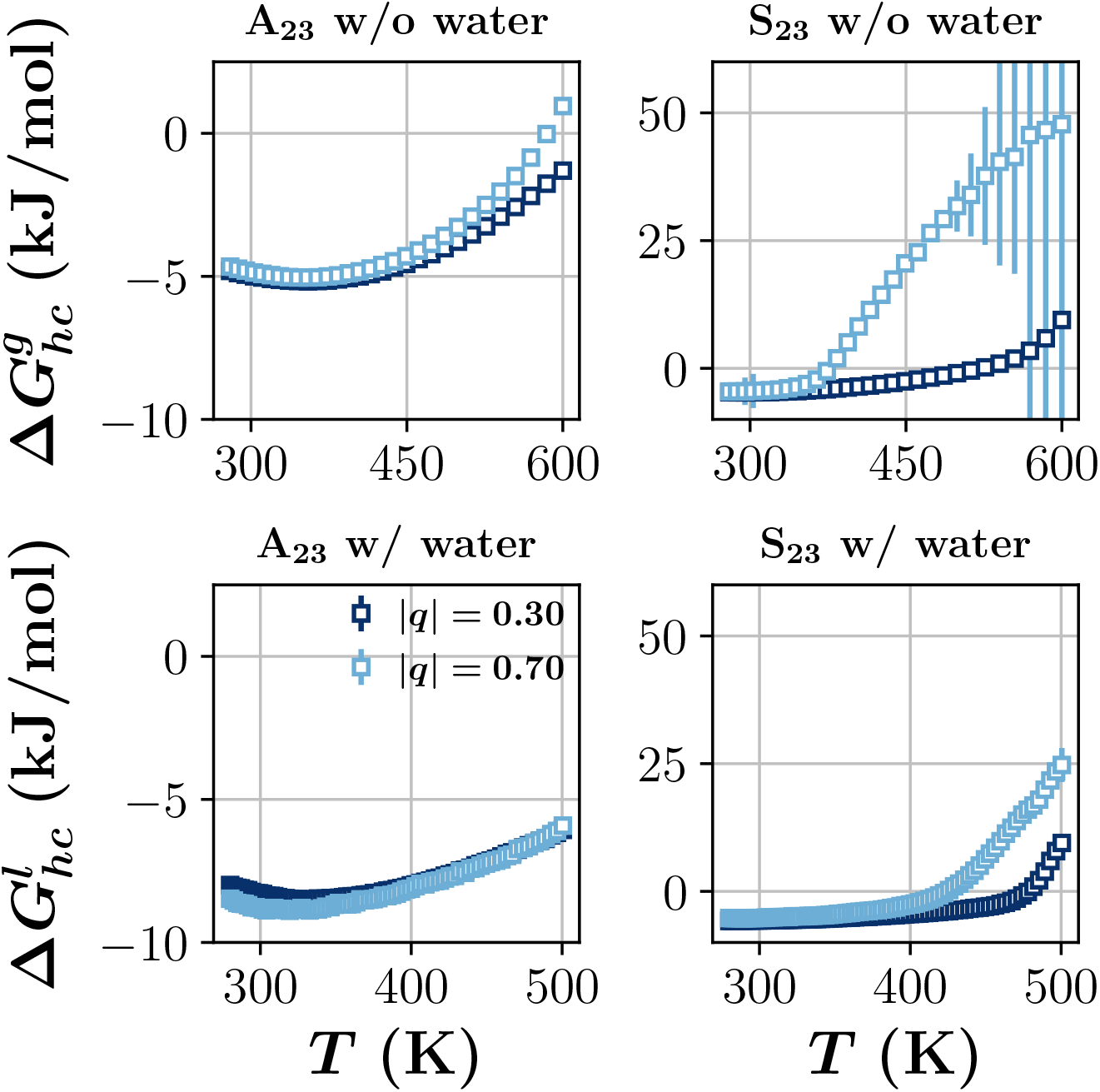
Free energy of helix formation in polyalanine and polyserine confined to a 13.6 Å boron nitride nanotube both with and without water. |*q*| is the absolute value of the partial charge placed on the boron and nitrogen atoms in the boron nitride nanotube.

### PNT/A_23_

The temperature dependence of *f_h_* is very similar in the PNT/A_23_ system compared to the NPNT/A_23_ system, with minor differences observed at a larger nanotube diameter of *D* = 14.9 Å (Fig. 2). The difference in the solvation free energies between the helix and coil states is approximately the same in both nanotubes with ΔΔ*G_hc_* ≈ −4kJ/mol (Table 2). However, there are disparities in the enthalpy and entropy differences in the two systems. For example, ΔΔ*H* is less negative in the PNT than the NPNT, but this is compensated by a positive ΔΔ*S* in the PNT suggesting the disparities in the thermodynamics arise from compensating water reorganization enthalpy and entropy. See the Supporting Information for further discussion. Thus, A_23_ appears agnostic to the nature of the nanotube surface, at least for a tube with relatively small partial charges without hydrogen-bonding groups. Surprisingly, the difference in free energies between helix and coil states ΔΔ*G_hc_* becomes more negative as the partial charge on the boron and nitrogen atoms increase inside the nanotube. This is likely because the water removed from the nanotube surface during helix formation in the |*q*| = 0.70 system has a larger entropy change 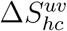 than the |*q*| = 0.30 system.

**Table 2:**
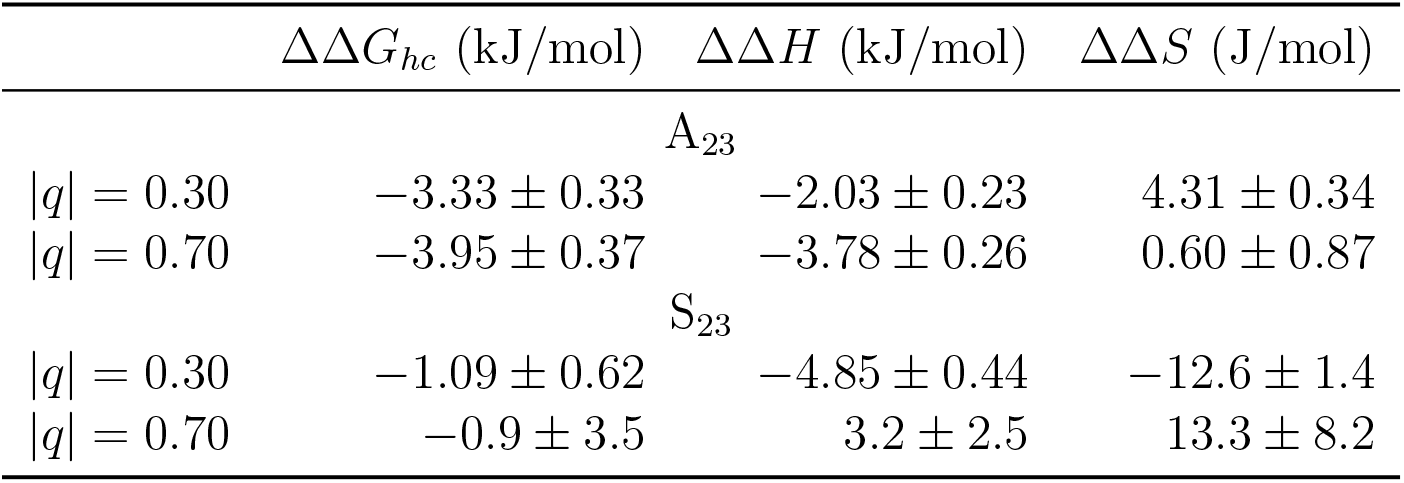
Differences in solvation free energy, enthalpy, and entropy for the helix and coil states at 300 K for A_23_ and S_23_ confined to the polar boron nitride nanotube.

### PNT/S_23_

Upon changing the nanotube walls from non-polar to polar, S_23_ forms a helix when confined to the *D* = 13.6 Å nanotube in the presence of water at 300 K. Further, the helix is more stable in the presence of water than in the gas phase with ΔΔ*G_hc_* ≈ −1 kJ/mol. The thermodynamics in the PNT (Table 2) demonstrate a dramatic reversal from S_23_ in the NPNT. Instead of a large positive enthalpy change as in the NPNT, the partial charge on the PNT has made it so ΔΔ*H* < 0. The electrostatic interaction between the peptide and nanotube mitigates the desolvation energy of the peptide upon helix formation (see the SI for further discussion), thereby stabilizing the helix state in the PNT. When |*q*| = 0.70 the uncertainty resulting from the fit to ln *K*(*T*) in the gas phase was too large to provide accurate estimates of the thermodynamics. With a slight increase in the nanotube diameter from *D* = 13.6 Å to *D* = 14.9 Å, the fraction of S_23_ forming a helix is nearly zero (Fig. 2) in the presence of water except in the nanotube with |*q*| = 0.70 partial charges on boron and nitrogen.

### NPNT/VSV-G

We also conducted gas phase and liquid water replica exchange MD simulations of wild type and three mutant amino acid sequences from the membrane protein VSV-G. The 20 amino acid wild type sequence was SSIASFFFIIGLIIGLFLVL. In the first mutant (AAIAAFFFIIGLIIGLFLVL) we substituted alanine for serine to observe what effect, if any, the substitution would have on the helix content of the protein sequence. In the second mutant (SSIASFFFIIALIIALFLVL) we substituted alanine for glycine since glycine is a known helix breaker in proteins. The third mutant contained both substitutions (AAIAAFFFIIALIIALFLVL). Figure 7 displays *f_h_* for each sequence as a function of nanotube diameter and protein sequence both with and without water while confined to the NPNT. In each case, the middle of the sequence forms a helix inside the nanotube with slight helicity towards the C-terminus of the sequence. This observation is consistent with results from molecular dynamics simulations of the same sequence confined to the ribosome tunnel. ^11^ The N-terminus of the sequence contains three serines that display little helicity inside the nanotube, and does not increase upon substitution with alanine. Confinement has the greatest effect on helix stabilization in alanine and serine at *D* = 13.6 Å while the amino acids with bulky side chains, such as isoleucine (I) and leucine (L), are too large to form a helix at that diameter. Therefore, we do not observe helix formation in alanine or serine for *D* ≥ 14.9 Å but we observe helix formation in the bulky amino acids in the middle of the protein sequence. These data highlight the extreme sensitivity of confinement induced helix stabilization on the diameter of the confining surface.

**Figure 7:**
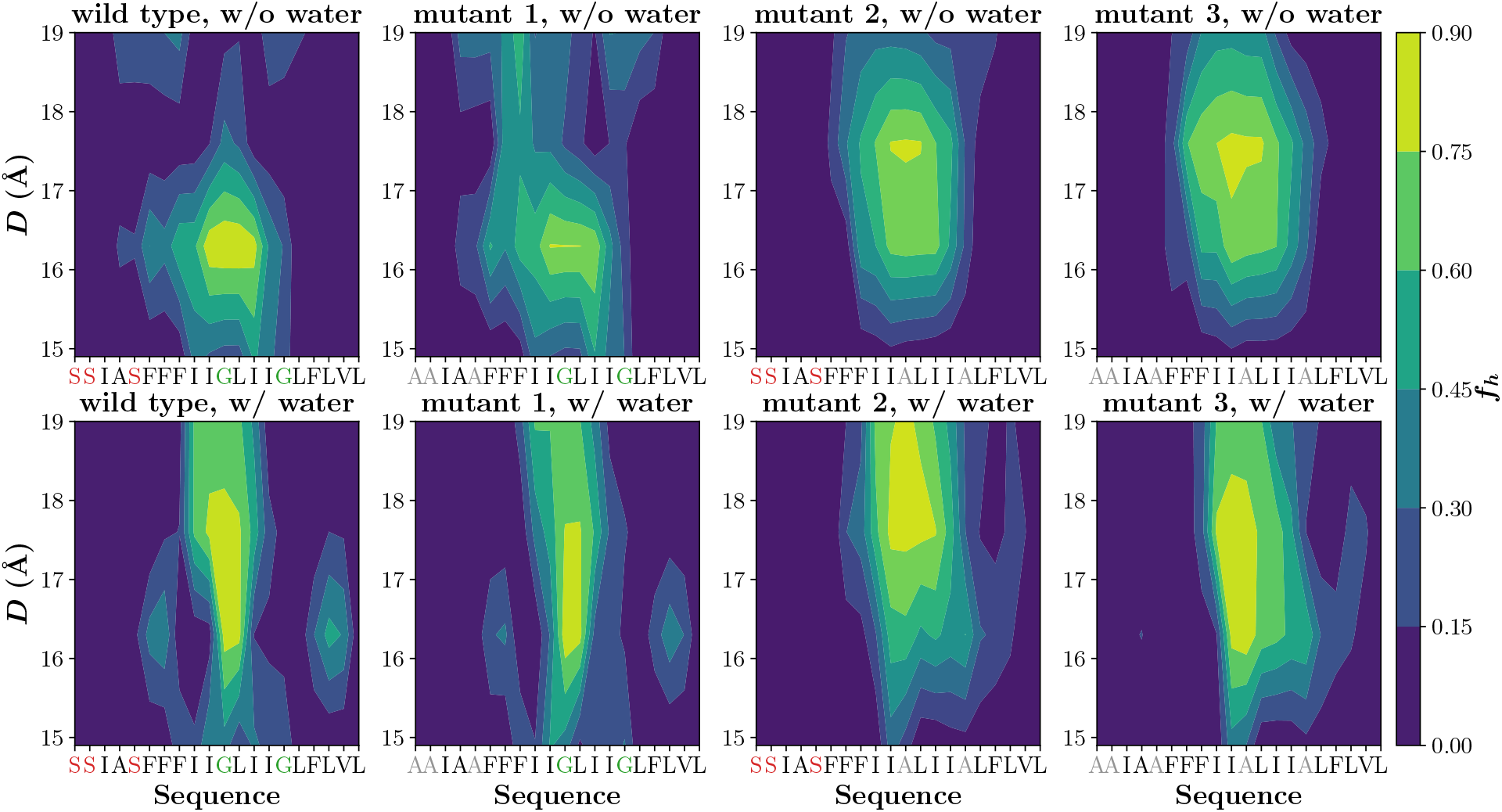
Displayed is the fractional helix content *f_h_* of wild-type and mutant sequences extracted from the protein VSV-G as a function of nanotube diameter and protein sequence. The top row is from simulations in the gas phase, while the simulations in the second row contained liquid water.

### Biological Implications

The results presented thus far demonstrate a complex interplay of many competing interactions contributing to the preferred states of proteins in nanotubes. Confinement to the nanotube in the gas phase induces helix formation when the diameter of the confining wall is slightly larger than the diameter of the helix (Fig. 2). This behavior is expected from prior studies utilizing elegant concepts from polymer physics. ^13^ However, even in the gas phase the extent of helix stabilization depends on the nature of the amino acid sequence and on the type of interactions with the nanotube walls. For example, polyalanine forms a helix inside both the non-polar and polar nanotubes, but polyserine displays differing behavior depending on the interactions with the nanotube, especially as the diameter of the nanotube increases (Fig. 2). In liquid water, water-mediated interactions between the protein and the nanotube further complicate the picture. For example, when the nanotube wall is non-polar, polyserine prefers the coil state. However, upon confinement to a nanotube with polar walls, the helix state is stabilized for diameters just slightly larger than the helix. These data suggest that protein sequences containing stretches of hydrophobic amino acids might preferentially form *α*-helices inside the ribosome tunnel. Indeed, experiments investigating helix formation inside the ribosome tunnel demonstrated that hydrophobic transmembrane protein sequences form compacted structures inside the tunnel, suggesting the formation of an *α*-helix.^11^

We performed simulations of a 20 amino acid sequence from wild-type and mutant forms of the VSV-G protein confined to a non-polar nanotube with varying diameters. In the wild-type and mutant sequences we observed helix formation in the middle of the sequence with some helicity at the C-terminus. The portion of the protein sequence containing serine did not form a helix inside the nanotube and this was expected when considering our results with S_23_. When alanine was substituted for serine in the sequence, the helicity did not increase which was contrary to our expectations arising from the A_23_ results. These observations can be rationalized when considering the strong diameter dependence of confinement induced helix stabilization. The smaller alanine does not experience confinement induced helix stabilization at the same diameters as the larger hydrophobic amino acids such as leucine (L), isoleucine (I), and phenylalanine (F). Consequently, the helicity does not increase upon substitution of serine with alanine for *D* ≥ 14.9 Å However, each VSV-G protein sequence formed a helix within the center of the sequence and this is consistent with prior studies. ^11^

## Concluding Remarks

In this paper, we present the results of replica exchange molecular dynamics to investigate the thermodynamics of helix-coil transitions of the polypeptides 23-alanine and 23-serine in open nanotubes. The transitions occur at diameters similar to the width of the ribosome tunnel, and are accompanied by the expulsion of water from the tubes when they are open to a water reservoir. We elucidate the effect of water by comparison with the corresponding results for systems in the gas phase. We also compare and contrast gas and liquid water phase replica exchange MD simulations of the wild type and mutant amino acid sequences from the membrane protein VSV-G.

Taken together, our results suggest that water-mediated interactions contribute to helix formation inside nanotubes. Several possibilities emerge for the water mediated interactions depending on the sequence of the peptides, which implies that it is difficult to construct a generic theory. However, we showed that helix formation is dependent on the difference in water interaction energy 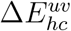 and entropy 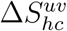 in the helix and coil states. Peptides containing stretches of hydrophobic amino acids preferentially form *α*-helices inside nanotubes, which is consistent with theories based on polymer physics. Our simulations show that protein sequences containing polar amino acids could also form *α*-helices but the extent of helix formation is sensitively dependent on the diameter of the confining nanotube and the nature of the nanotube surface. The different possibilities illustrated here also are consistent with experiments that show that structure formation in the ribosome tunnel depends on sequence. Our work shows that these variations are caused by water-mediated interactions, which implies that predicting the stability of proteins under confinement (in the cavity of GroEL or peptides in the ribosome tunnel) will require accounting for the effects of water. In this context, coarse-grained models incorporating water-mediated interactions^59,60^ may prove beneficial.

## Acknowledgement

We thank Stephen Cousins of the University of Maine Advanced Computing Group for significant allotments of computing time and his technical assistance. This work was supported by the National Science Foundation (CHE 19-00093) and the Welch Foundation through grant F-0019 administered through the Collie-Welch chair.

## Supporting Information Available

Additional details about the simulation methods and analysis, thermodynamics of helix formation for each system described in the main text, and interaction energies as a function of time and their relation to the solvation thermodynamics.

## Graphical TOC Entry

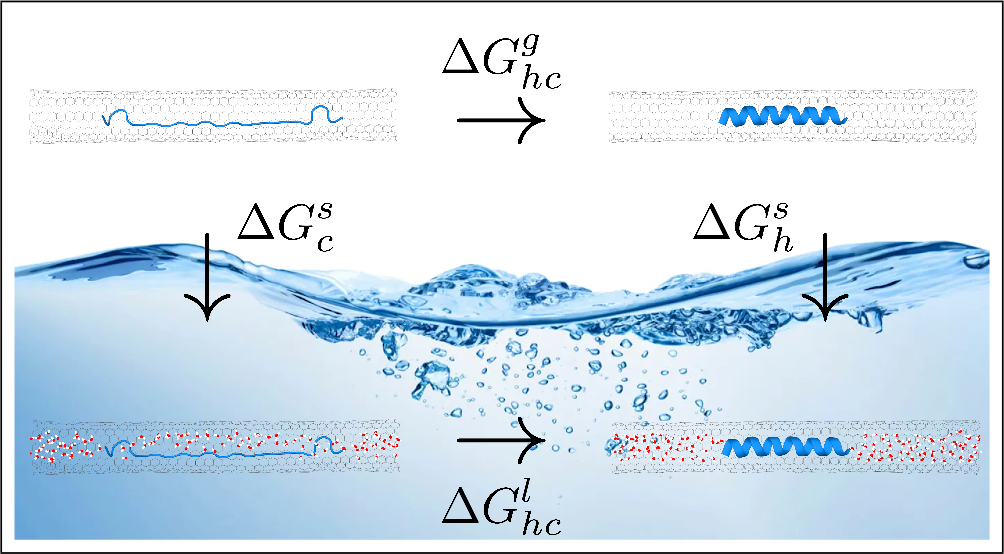

