## Supplemental Data for "Water-mediated interactions determine helix formation of peptides in open nanotubes"

### Supporting Information for Water-mediated interactions determine helix formation of peptides in open nanotubes

This Supporting Information (SI) contains supporting information relating to the simulation and analysis methods. We also provide results for the thermodynamics of the helix-coil transition for $A_{23}$ and $S_{23}$ confined within the polar and non-polar nanotubes.

### Methods

#### Simulation Details

The systems were constructed by placing a capped 23-alanine ($CH_3CO$-$A_{23}$-$NHCH_3$) or 23-serine ($CH_3CO$-$S_{23}$-$NHCH_3$) polypeptide inside the nanotubes coincident with the axis running through the center of the tube. The alanine polypeptide started from a fully extended ($\psi = 180°, \phi = -180°$) conformation while the serine polypepide started from the $\alpha$-helix state. The nanotube and polypeptide were hydrated in a TIP3P water bath with hexagonal periodic boundary conditions. During the hydration procedure water was allowed inside the CNT. Water molecules within 3 Å of the serine polypetide were removed following solvation. The purpose of this was to avoid starting the simulation in a physically unrealistic high energy state. The system was energy minimized using the conjugate gradient and steepest descent method. We used GROMACS 5.1.2 to perform the simulations with a time step of 0.002 ps in conjunction with constrained bonds to hydrogen atoms with the Lincs algorithm. Equilibration was performed in the NPT ensemble for 1 ns with Langevin dynamics and isotropic Berendsen pressure coupling at 1 bar. The inverse friction constant in the Langevin algorithm was taken to be 2 ps. This results in a friction that is lower than that of water but high enough to remove excess heat. Langevin dynamics was chosen as a thermostat because it has been shown to reliably preserve the canonical ensemble in replica exchange simulations.[1] The time constant for the Berendsen pressure coupling was 1 ps. In the liquid water simulations, non-bonded interactions were evaluated with a cutoff of 10 Å. A long range dispersion was calculated for the energy and pressure. The particle mesh Ewald (PME) method was used to evaluate the electrostatics with a grid spacing of 1.2 Å. The non-bonded interactions were evaluated with no cut-off in the gas phase simulations. The temperature spacing for the 30 replicas spanning the temperature range 280 K to 600 K in the gas phase was determined by following a procedure established elsewhere.[2] The CNTs were held fixed by restraining the center of mass (COM) coordinate of 8 carbon atoms in the middle of the CNT. The COM coordinate was scaled with the scaling matrix of the pressure coupling.

#### Boron Nitride Nanotube

Simulation parameters for the boron nitride nanotube were taken from a previous study.[3–5]

#### Analysis

As with our previous work,[6] we monitored the fractional helix content of the polypeptides with time to ensure we were sampling from equilibrium structures. We performed a time series analysis of the fractional helix content where we calculated the autocorrelation of $f_h$ and the effective number of uncorrelated samples and the statistical inefficiency.[7] We also checked that the choice of algorithm was providing simulation results from the correct thermodynamic ensemble.[8] Thus, we are confident that our simulation data is representative of equilibrium structures from valid thermodynamic ensembles.

The thermodynamics at 300 K were determined by fitting $\Delta G_{hc} = -RT \ln K(T)$ to the following function,

$$\begin{aligned}\Delta G_{hc} = H_{300\,\mathrm{K}} &+ a(T^2 - (300\,\mathrm{K})^2)/2 + b(T^3 - (300\,\mathrm{K})^3)/3 + c(T^4 - (300\,\mathrm{K})^4)/4 \\ &+ d(T^5 - (300\,\mathrm{K})^5)/5 + e(T^6 - (300\,\mathrm{K})^6)/6 + f(T^7 - (300\,\mathrm{K})^7)/7 \\ &+ g(T^8 - (300\,\mathrm{K})^8)/8 - T\{S_{300\,\mathrm{K}} + a(T - 300\,\mathrm{K}) + b(T^2 - (300\,\mathrm{K})^2)/2 \\ &+ c(T^3 - (300\,\mathrm{K})^3)/3 + d(T^4 - (300\,\mathrm{K})^4)/4 + e(T^5 - (300\,\mathrm{K})^5)/5 \\ &+ f(T^6 - (300\,\mathrm{K})^6)/6 + g(T^7 - (300\,\mathrm{K})^7)/7\} \end{aligned} \tag{1}$$

where $a$, $b$, $c$, $d$, $e$, $f$, and $g$ are fitting parameters. The fit was performed for each system with an increasing number of fitting parameters until the thermodynamic values were observed not to vary more than a tenth between fits. The following table lists the number of fitting parameters that were used for each system. For the $S_{23}$ polypeptide in the NPNT in liquid water we used the following equation to determine the thermodynamics,

$$\Delta G_{hc} = H_{300\,\mathrm{K}} + C_p(T - 300\,\mathrm{K}) - T(S_{300\,\mathrm{K}} + C_p \ln[\frac{T}{300\,\mathrm{K}}]) \quad (2)$$

where $C_p$ is the constant pressure heat capacity.

**Table 1: Number of fitting parameters from Eq. (1) for each system. *See Eq. (2) for the equation used in the fitting procedure in this case.**

| system | # of parameters |
|---|---|
| $A_{23}$, NPNT, water | 4 |
| $A_{23}$, NPNT, no water | 4 |
| $A_{23}$, PNT, water | 4 |
| $A_{23}$, PNT, no water | 4 |
| $S_{23}$, NPNT, water | * |
| $S_{23}$, NPNT, no water | 7 |
| $S_{23}$, PNT, water | 5 |
| $S_{23}$, PNT, no water | 7 |

### Results

In the following we provide data for thermodynamics of the helix-coil transition in liquid and gas phases. We also discuss the interaction energies of the $A_{23}$/NPNT, $A_{23}$/PNT, $S_{23}$/NPNT, and $S_{23}$/PNT systems and relate them to the discussion of the solvation thermodynamics discussed in the main text.

#### Thermodynamics in the liquid and gas phases

**Table 2: Free energy, enthalpy, and entropy of helix formation at 300 K inside the 13.6 Å carbon nanotube both with and without water.**

| | $\Delta G_{hc}$ (kJ/mol) | $\Delta H$ (kJ/mol) | $\Delta S$ (J/mol) |
|---|---|---|---|
| | $A_{23}$ w/o water | | |
| $\lambda = 1.00$ | $-4.97 \pm 0.07$ | $-2.94 \pm 0.05$ | $6.76 \pm 0.16$ |
| $\lambda = 0.56$ | $-5.20 \pm 0.08$ | $-3.46 \pm 0.05$ | $5.81 \pm 0.18$ |
| | $S_{23}$ w/o water | | |
| $\lambda = 1.00$ | $-4.35 \pm 0.77$ | $-4.72 \pm 0.55$ | $-1.2 \pm 1.8$ |
| $\lambda = 0.56$ | $-4.44 \pm 0.66$ | $-5.79 \pm 0.46$ | $-4.5 \pm 1.5$ |
| | $A_{23}$ w/ water | | |
| $\lambda = 1.00$ | $-8.75 \pm 0.24$ | $-7.35 \pm 0.17$ | $4.67 \pm 0.56$ |
| $\lambda = 0.56$ | $-9.62 \pm 0.38$ | $-12.75 \pm 0.27$ | $-10.43 \pm 0.89$ |
| | $S_{23}$ w/ water | | |
| $\lambda = 1.00$ | $6.7 \pm 2.2$ | $44.1 \pm 1.6$ | $124.4 \pm 4.9$ |
| $\lambda = 0.56$ | $4.8 \pm 1.4$ | $20.0 \pm 1.0$ | $50.8 \pm 3.1$ |

**Table 3: Free energy, enthalpy, and entropy of helix formation in $\mathbf{A_{23}}$ and $\mathbf{S_{23}}$ at 300 K inside the 13.6 Å polar boron nitride nanotube both with and without water.**

| | $\Delta G_{hc}$ (kJ/mol) | $\Delta H$ (kJ/mol) | $\Delta S$ (J/mol) |
|---|---|---|---|
| | $A_{23}$ w/o water | | |
| $\lvert q\rvert = 0.30$ | $-4.97 \pm 0.05$ | $-2.86 \pm 0.03$ | $7.03 \pm 0.11$ |
| $\lvert q\rvert = 0.70$ | $-4.83 \pm 0.10$ | $-2.60 \pm 0.07$ | $7.45 \pm 0.23$ |
| | $S_{23}$ w/o water | | |
| $\lvert q\rvert = 0.30$ | $-4.66 \pm 0.33$ | $-3.56 \pm 0.23$ | $3.69 \pm 0.78$ |
| $\lvert q\rvert = 0.70$ | $-4.3 \pm 3.5$ | $-9.1 \pm 2.5$ | $-16.4 \pm 8.1$ |
| | $A_{23}$ w/ water | | |
| $\lvert q\rvert = 0.30$ | $-8.30 \pm 0.13$ | $-4.90 \pm 0.10$ | $11.35 \pm 0.31$ |
| $\lvert q\rvert = 0.70$ | $-8.78 \pm 0.36$ | $-6.37 \pm 0.25$ | $8.05 \pm 0.84$ |
| | $S_{23}$ w/ water | | |
| $\lvert q\rvert = 0.30$ | $-5.75 \pm 0.52$ | $-8.41 \pm 0.37$ | $-8.9 \pm 1.2$ |
| $\lvert q\rvert = 0.70$ | $-5.16 \pm 0.46$ | $-5.96 \pm 0.33$ | $-2.7 \pm 1.1$ |

#### $A_{23}$/NPNT Energies

$A_{23}$ starts in the coil state at $t = 0$ and proceeds to form a helix in less than 50 ns in the $D = 13.6$ Å nanotube at 300 K as displayed in Fig. 1. Concomitantly, the peptide-water interaction energy increases with time, as does the nanotube-water dispersion energy (Fig. 2 (a) and (b)), but the water-water interaction energy decreases with time (Fig. 2 (c)). As discussed in the main text, the final result is an increase in the solute-solvent interaction energy in going from the coil to the helix state with the peptide-water Coulomb interaction energy making the largest contribution to the change. The water-water interaction energy decreases during helix formation. Therefore, $\Delta E_{uv} > 0$ and $\Delta\Delta H_{vv} < 0$ when $A_{23}$ transitions from the coil to the helix.

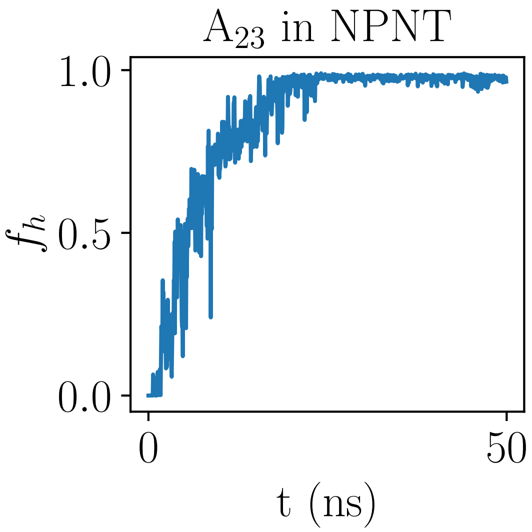


Figure 1: Fraction helix $f_h$ of $A_{23}$ inside the non-polar carbon nanotube as a function of time. Each data point is an average over 50 ps. At $t = 0$ the polypeptide is in the coil state.

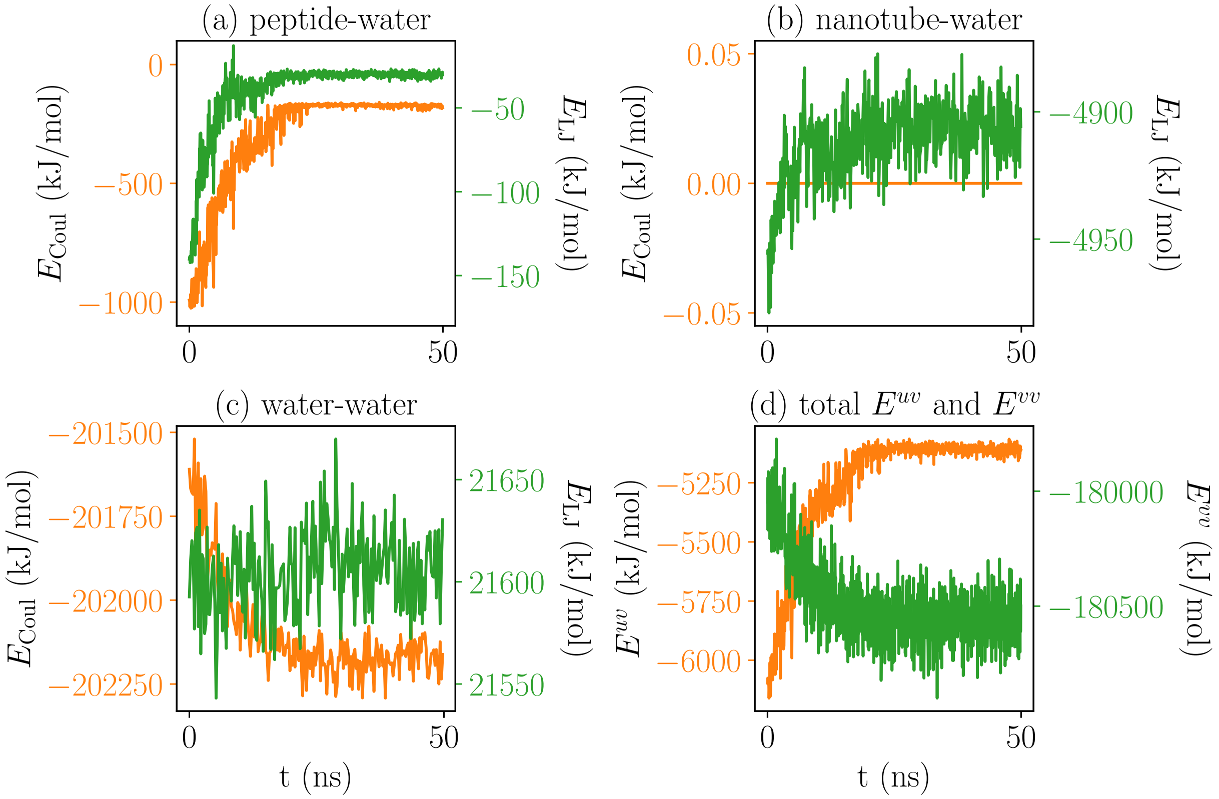


Figure 2: Energies for $A_{23}$ inside the $D = 13.6$ Å non-polar carbon nanotube as a function of time at 300 K. The short-range Coulomb $E_{\mathrm{Coul}}$ (orange) and dispersion $E_{\mathrm{LJ}}$ (green) energy for (a) peptide-water, (b) nanotube-water, and (c) water-water interactions. (d) Comparison of the total interaction energy for the solute (polypeptide/nanotube)-solvent (water) $E^{uv}$ (orange) and water-water $E^{vv}$ (green) interactions.

#### $S_{23}$/NPNT Energies

The folding time for $S_{23}$ was longer than a simulation time of 250 ns/replica. Consequently, data was acquired for the unfolding of $S_{23}$ from the helix to the coil. Figure 3 displays the fraction helix as a function of time at 300 K inside the $D = 13.6$ Å non-polar carbon nanotube. As can be seen from Fig. 3, the helix unfolds very rapidly—in less than 1 ns. Figure 4 indicates that the peptide-water and nanotube-water interaction energies decrease during unfolding, and the water-water interaction energy increases. Therefore, $\Delta E^{uv} > 0$ and $\Delta\Delta H^{vv} < 0$ for the coil to helix transition. However, in contrast to $A_{23}$, $S_{23}$ forms additional hydrogen bonds with water as indicated by $E_{\text{Coul}} < 1500$ kJ/mol in Fig. 4 (a) for the coil state, whereas $E_{\text{Coul}} \approx 1000$ kJ/mol in the coil state for $A_{23}$. Therefore, $\Delta E^{uv} >> 0$ for $S_{23}$ and this opposes helix formation.

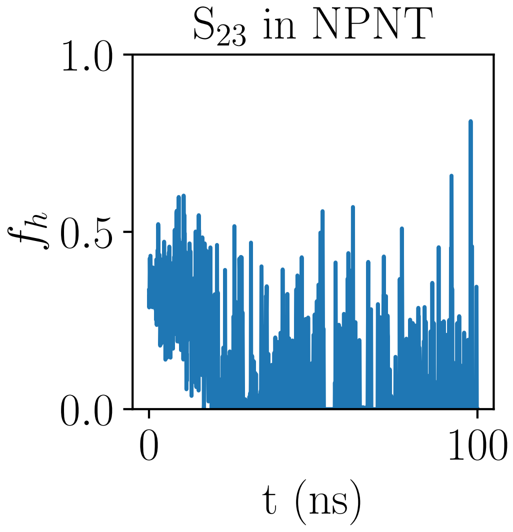


Figure 3: Fraction helix $f_h$ of $S_{23}$ inside the non-polar carbon nanotube as a function of time. Each data point is an average over 50 ps. At $t = 0$ the polypeptide is in the helix state.

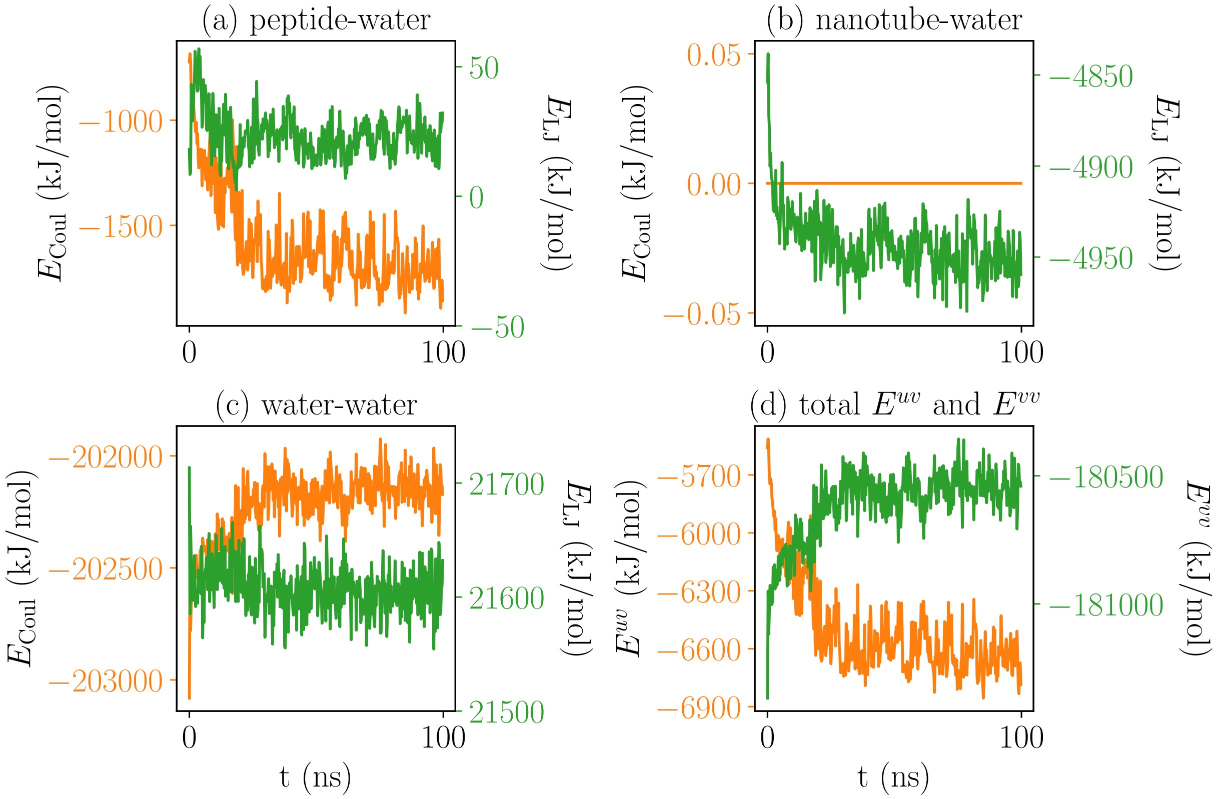


Figure 4: Energies for $S_{23}$ inside the $D = 13.6$ Å non-polar carbon nanotube as a function of time at 300 K. The short-range Coulomb $E_{\text{Coul}}$ (orange) and dispersion $E_{\text{LJ}}$ (green) energy for (a) peptide-water, (b) nanotube-water, and (c) water-water interactions. (d) Comparison of the total interaction energy for the solute (polypeptide/nanotube)-solvent (water) $E^{uv}$ (orange) and water-water $E^{vv}$ (green) interactions.

#### $A_{23}$/PNT Energies

The polar boron nitride nanotube has partial charges on the boron and nitrogen atoms which contribute to the interaction energy with water (Fig. 8 (b)). Further, the boron and nitrogen atoms have different dispersion energies than the carbon atoms in the CNT. These differences shift $E^{uv}$ compared to the $A_{23}$/NPNT system. However, $\Delta E^{uv}$ is mostly determined by the peptide-water interactions. Thus, the free energy of helix formation of $A_{23}$ inside the PNT is not much different than in the NPNT. The water-water interaction energy is also affected by the change in the nature of the nanotube, but because $\Delta H^{vv} = T\Delta S^{vv}$, the altered water-water interaction energy does not contribute to free energy of helix formation. Thus, $A_{23}$ appears agnostic to the nature of the nanotube surface, at least for relatively small partial charges on the nanotube atoms.

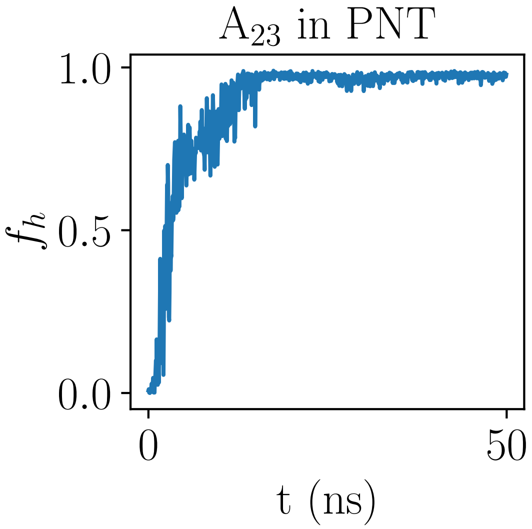


Figure 5: Fraction helix $f_h$ of $A_{23}$ inside the polar boron nitride nanotube as a function of time. Each data point is an average over 50 ps. At $t = 0$ the polypeptide is in the coil state.

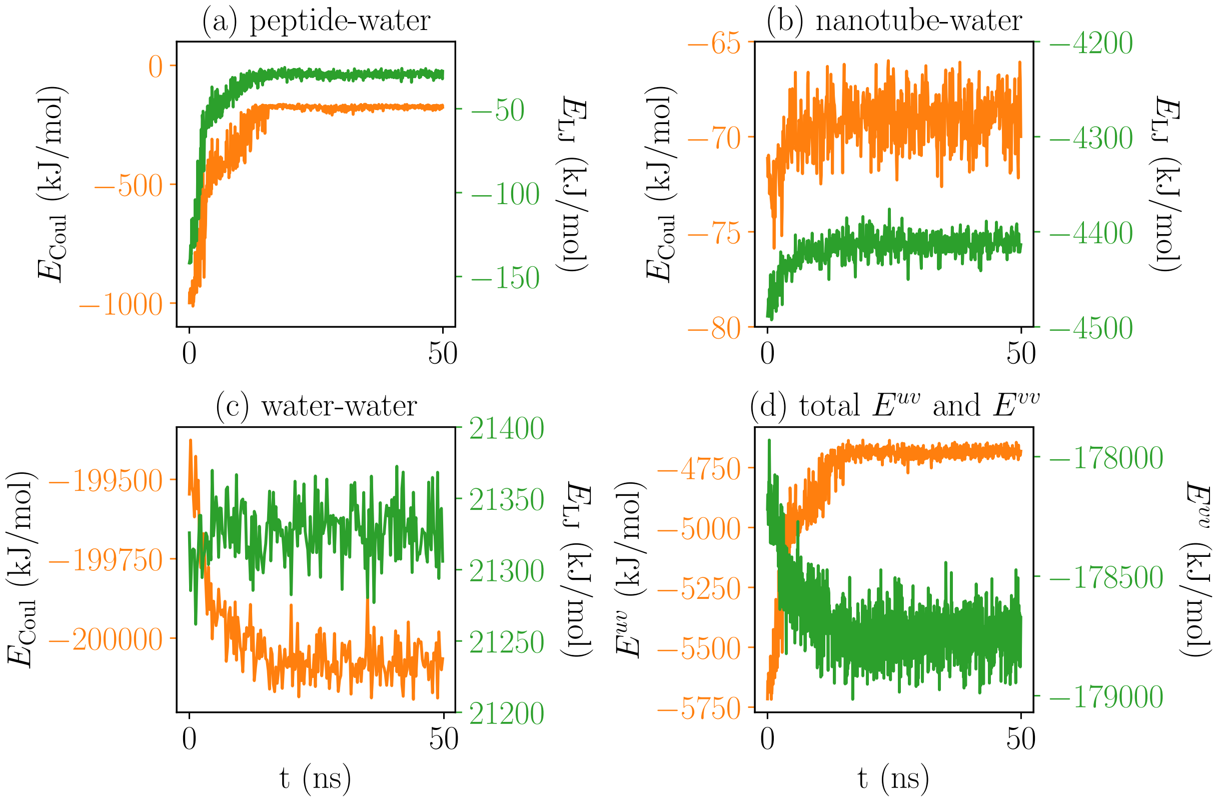


Figure 6: Energies for $A_{23}$ inside the polar boron nitride nanotube as a function of time. The short-range Coulomb $E_{\mathrm{Coul}}$ (orange) and dispersion $E_{\mathrm{LJ}}$ (green) energy for (a) peptide-water, (b) nanotube-water, and (c) water-water interactions. (d) Comparison of the total interaction energy for the solute (polypeptide/nanotube)-solvent (water) $E^{uv}$ (orange) and water-water $E^{vv}$ (green) interactions.

#### $S_{23}$/PNT Energies

In the case of $S_{23}$ confined to the PNT, the Coulomb interaction energy between $S_{23}$ and the nanotube (Fig. 9) is sufficient to offset the energy that would be gained if the peptide were to unfold and the side chains hydrated. Therefore, the electrostatic interaction with the nanotube stabilizes the helix state for the polar $S_{23}$ peptide.

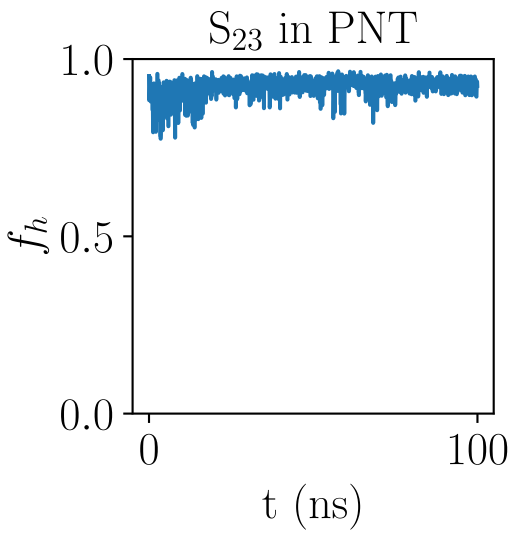


Figure 7: Fraction helix $f_h$ of $S_{23}$ inside the polar boron nitride nanotube as a function of time. Each data point is an average over 50 ps. At $t = 0$ the polypeptide is in the helix state.

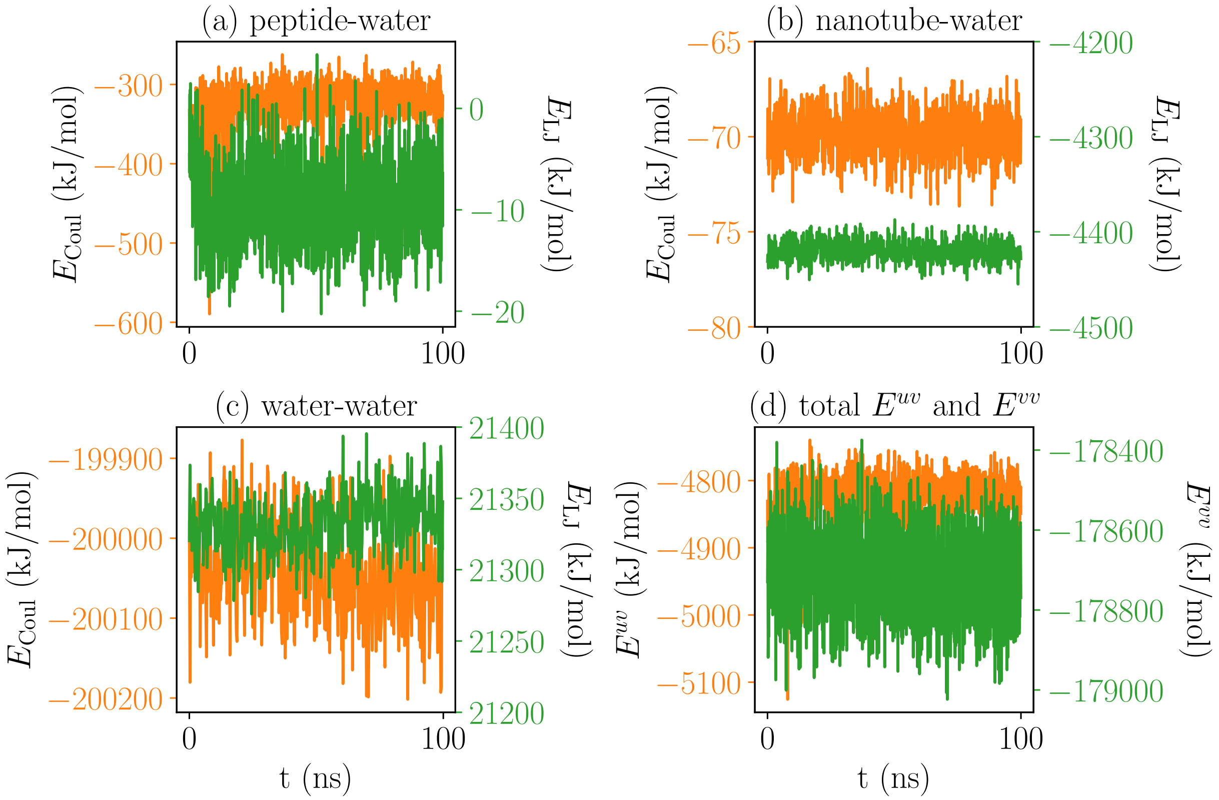


Figure 8: Energies for $S_{23}$ inside the polar boron nitride nanotube as a function of time. The short-range Coulomb $E_{\mathrm{Coul}}$ (orange) and dispersion $E_{\mathrm{LJ}}$ (green) energy for (a) peptide-water, (b) nanotube-water, and (c) water-water interactions. (d) Comparison of the total interaction energy for the solute (polypeptide/nanotube)-solvent (water) $E^{uv}$ (orange) and water-water $E^{vv}$ (green) interactions.

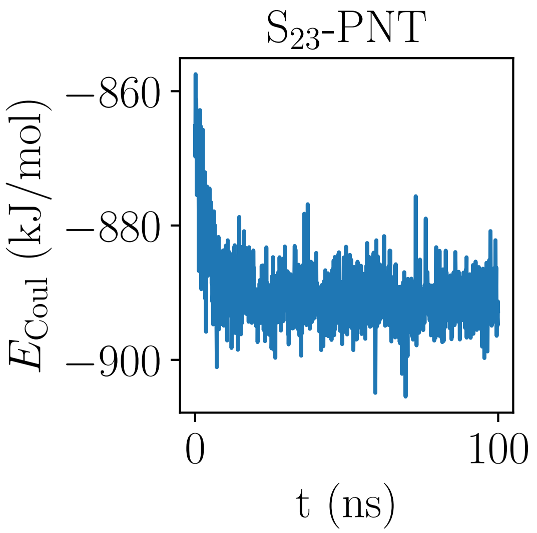


Figure 9: Coulomb interaction energy between $S_{23}$ and the polar boron nitride nanotube as a function of time. Each data point is an average over 50 ps. At $t = 0$ the polypeptide is in the helix state.
